# Particle-templated emulsification for microfluidics-free digital biology

**DOI:** 10.1101/304923

**Authors:** Makiko N. Hatori, Samuel C. Kim, Adam R. Abate

## Abstract

The compartmentalization of reactions in monodispersed droplets is valuable for applications across biology. However, the requirement of microfluidics to partition the sample into monodispersed droplets is a significant barrier that impedes implementation. Here, we introduce particle-templated emulsification, a method to encapsulate samples in monodispersed emulsions without microfluidics. By vortexing a mixture of hydrogel particles and sample solution, we encapsulate the sample in monodispersed emulsions that are useful for most droplet applications. We illustrate the method with ddPCR and single cell culture. The ability to encapsulate samples in monodispersed droplets without microfluidics should facilitate the implementation of compartmentalized reactions in biology.

## Introduction

Biological systems are often complex, necessitating many experiments to characterize them ^1–3^. The modern biological lab is thus full of high-throughput tools, from robotic fluid handlers that automate experiments to flow cytometers, next generation sequencers, and mass spectrometers for high-throughput sample analysis ^4,5^. Nevertheless, the complexity of biology outpaces even these instruments, motivating new tools of ever higher throughput ^6^. In particular, microfluidic-based single-cell sequencing has become a powerful tool to study cell populations and heterogeneity^7–9^.

The field of droplet microfluidics studies how laboratory automation can be scaled up by reducing reaction volumes to picoliters and increasing processing speed to kilohertz ^10–13^. Microfluidic devices form, process, and sort aqueous droplets suspended in carrier oil, each acting as an isolated “test tube” in which a reaction can be performed with a single molecule or a cell^7,8,14^. The throughput of the approach, combined with the tiny reagent consumption, provide unprecedented potential for a new era of high-throughput biology. Indeed, droplet methods have already been applied to numerous important applications, including enzyme evolution, single cell sequencing, and accurate DNA and protein quantitation ^7,15–17^.

A barrier to implementing powerful droplet methods is the requirement for specialized microfluidic hardware and skill. Consequently, the most notable impacts have been made when a specific system is commercialized and reduced to a user-friendly instrument, such as digital droplet PCR (ddPCR) and ultrahigh-throughput single cell transcriptome sequencing ^18,19^. Nevertheless, engineers have developed many other microfluidic tools that have yet to achieve the high bar of commercialization. The inability to immediately translate technological advances in droplet microfluidics to the biology lab is thus a major inefficiency in microfluidics research. To address this bottleneck, a new strategy for performing isolated reactions in uniform droplets without microfluidics is needed.

In this paper, we describe particle-templated emulsification (PTE), an approach for generating compartmentalized reactions in monodispersed droplets with vortexing. The central concept of PTE is to use particles to “template” the formation of droplets of similar size under agitation with oil. To encapsulate a sample into monodispersed droplets, as is normally accomplished with microfluidics, monodispersed particles are used. If cells, DNA molecules, or beads are present in the sample, they are co-encapsulated with the particles. The droplet-templating particles can also introduce additional components to the droplets, such as barcoded oligonucleotides required for single cell transcriptome sequencing, or reagents necessary for cell lysis or reporter assays ^7,20^. To demonstrate the power of PTE for microfluidics-free digital biology, we use it to perform ddPCR and single-cell culture.

## Experimental section

### Hydrogel particle preparation

6.2% acrylamide (Sigma-Aldrich), 0.18% N,N’-Methylenebisacrylamide (Sigma-Aldrich) and 0.3% ammonium persulphate (Sigma-Aldrich) is used for PAA particle generation. 14%(w/v) 8-arm PEGSH (Creative PEGworks) in 100 mM NaHCO_3_ and PEGDA (6 kDa, Creative PEGworks) in 100 mM NaHCO_3_ are used for PEG particle generation. 1% low melting temperature agarose (Sigma-Aldrich) is used for agarose particle generation. The agarose solution is warmed with a space heater during emulsification to prevent solidification. Agarose and PEG solutions are injected into a droplet generation device (Supplementary Fig. S2) with the oil (HFE-7500 fluorinated oil supplemented with 5%(w/w) deprotonated Krytox 157 FSH) using syringe pumps (New Era, NE-501). The PAA solution is injected into the droplet generation device with the fluorinated oil supplemented with 1% TEMED. The hydrogel solution and oil are loaded into separate 1-mL syringes(BD) and injected at 300 and 500 μl, respectively, into the droplet generation device using syringe pumps, controlled with a custom Python script (https://github.com/AbateLab/Pump-Control-Program). The PAA and PEG droplets are collected and incubated for one hour at room temperature for gelation. The agarose droplets are incubated on ice for gelation. After gelation, the gelled droplets are transferred to an aqueous carrier by destabilizing them in oil with addition of an equal volume of 20%(v/v) perfluoro-1-octanol in HFE-7500. The particles are washed twice with hexane containing 2% Span-80 (Sigma-Aldrich) to remove residual oil. Following the hexane wash, the particles are washed with sterile water until all oil is removed. Droplets are imaged using the EVOS Cell Imaging System (Thermo Fisher). Images are taken under a 4x and 10x objective using EVOS FITC LED light sources.

### Device fabrication

The polydimethylsiloxane (PDMS) device used for making monodispersed hydrogel particles is fabricated by pouring uncured PDMS (10:1 polymer-to-crosslinker ratio) over a photolithographically patterned layer of photoresist (SU-8 3025, MicroChem) on a silicon wafer. The device is cured for one hour in an 80°C oven, excised with a scalpel and inlet ports are punched using a 0.75-mm biopsy puncher (World Precision Instruments, *#*504529). The device is bonded to a glass slide using oxygen plasma and the inner surface of the channels treated with Aquapel (PPG Industries) to render them hydrophobic. The sealed device is baked at 80°C for 10 min.

### ddPCR

Monodispersed PAA particles or commercial PAA particles (Bio-Rad, 150-4164) are washed with 0.5% Triton-X100 (Sigma-Aldrich) in sterile water. 33 *μ*L of washed PAA particles are mixed with 17 *μ*L of PCR reagents to make a total volume of 50 *μ*L. The 50 *μ*L mixture includes 1x LongAmp Taq Reaction Buffer (NEB), 2 units of LongAmp Taq DNA Polymerase (NEB), 0.6 *μ*M of forward and reverse primers (IDT), 0.6 *μ*M of TaqMan probe (IDT), 300 *μ*M of dNTPs (Fisher Scientific) and a varying amount of budding yeast *Saccharomyces cerevisiae* genomic DNA (Milipore). For the multiplexed ddPCR, additional 0.6 *μ*M of forward and reverse primers and TaqMan probe for lambda virus DNA are included. The primer/probe sequences are as follows:

Yeast FWD 5’-GCAGACCAGACCAGAACAAA-3’,
Yeast REV 5’-ACACGTATGTATCTAGCCGAATAAC-3’,
Yeast Probe 5’-/56-FAM/ATATGTTGT/ZEN/TCACTCGCGCCTGGG/3IABkFQ/-3’,
Lambda FWD 5’-GTGGCATTGCAGCAGATTAAG-3’,
Lambda REV 5’-GGC AGTGAAGCCCAGATATT-3’,
Lambda Probe 5’-/Cy5/TATCCGTCAGGCAATCGACCGT G/3IAbRQSp/-3’.

The mixture is incubated for 15 min to allow PCR reagent to diffuse into the particles, and centrifuged for 1 min at 6,000 g. Excess aqueous phase is removed with a micropipette. 20 *μ*L of particles and 25 *μ*L of HFE-7500 oil supplemented with 2% (w/w) PEG-PFPE amphiphilic block copolymer surfactant (008-Fluoro-surfactant, RAN Biotechnologies) are mixed well by tapping in a 1.7-mL Eppendorf tube. The mixture is agitated at 2,300 rpm for 30 sec with a vortex (VWR). After transferring the emulsion to PCR tubes, the oil under the buoyant droplets is removed with a pipette and replaced with FC-40 oil (Sigma-Aldrich) containing 5% (w/w) PEG-PFPE amphiphilic block copolymer surfactant. This oil/surfactant combination yields greater thermostability during PCR. The emulsion is transferred to a T100 thermocycler (Bio-Rad) and subjected to the following program: 94°C for 30 sec, followed by 35 cycles of 94°C for 30 sec, 53°C for 60 sec and 65°C for 50 sec, followed by final extension of 10 min at 65°C and held at 12°C. The droplets are imaged using the EVOS Cell Imaging System (Thermo Fisher) under a 10x and 20x objective with EVOS GFP and FITC LED light sources.

### Cell culture

A budding yeast *Saccharomyces cerevisiae* strain expressing yellow fluorescent protein (YFP) is grown at 30°C in a standard rich media (YPD) and the cell density measured using NanoDrop (Thermo Fisher). PAA particles are washed with 0.5% Triton in YPD and centrifuged to remove excess aqueous solution. Diluted yeast suspension is prepared to achieve Poisson-distributed cell occupancy per droplet. 1 *μ*L of yeast suspension, 20 *μ*L of particles and 25 *μ*L of HFE-7500 oil with 2% (w/w) PEG-PFPE amphiphilic block copolymer surfactant are mixed well by tapping in a 1.7-mL Eppendorf tube. The mixture is vortexed at 2,300 rpm for 30 sec. Holes are punched on the tube lid for oxygen exchange for yeast cells and 1 mL of YPD media added. Cells are incubated at 30°C for 10 hours and imaged using the EVOS Cell Imaging System under 20x objective with EVOS RFP and FITC LED light sources.

## Results and Discussion

The objective of PTE is to use vortexing to encapsulate biological samples in uniform droplets, which have been shown by the microfluidics community to be valuable for many applications ^21,22^. The primary challenge is that, while vortexing is simple, uncontrolled agitation normally generates a broad distribution of droplet sizes ^23^. The strategy of PTE is to obtain a defined droplet size by using monodispersed particles to “template” the formation of monodispersed droplets. Hydrogel particles comprising >95% aqueous solution are used for the templating, so that the final droplet is mostly aqueous solution, necessary for performing biochemical reactions in the resultant droplets. The particles are added to the solution to be encapsulated, with oil and surfactant, and the mixture vortexed. The hydrogel particles are permeable to molecules with hydraulic diameters smaller than the pore size, like small molecules, but are impermeable to large molecules, such as genomic DNA, which remain within the thin layer of aqueous solution surrounding the particles in the droplets^22^ (Fig. 1a-e). During vortexing, the particles are dispersed into continually smaller droplets until each contains just one particle and a thin shell of aqueous solution, as illustrated in (Fig. 1e). Beyond this, further droplet breakup is suppressed because it requires fracturing solid particles. The result is an emulsion in which the droplets are of a similar size to the original particles and, thus, monodispersed.

**Figure 1.**
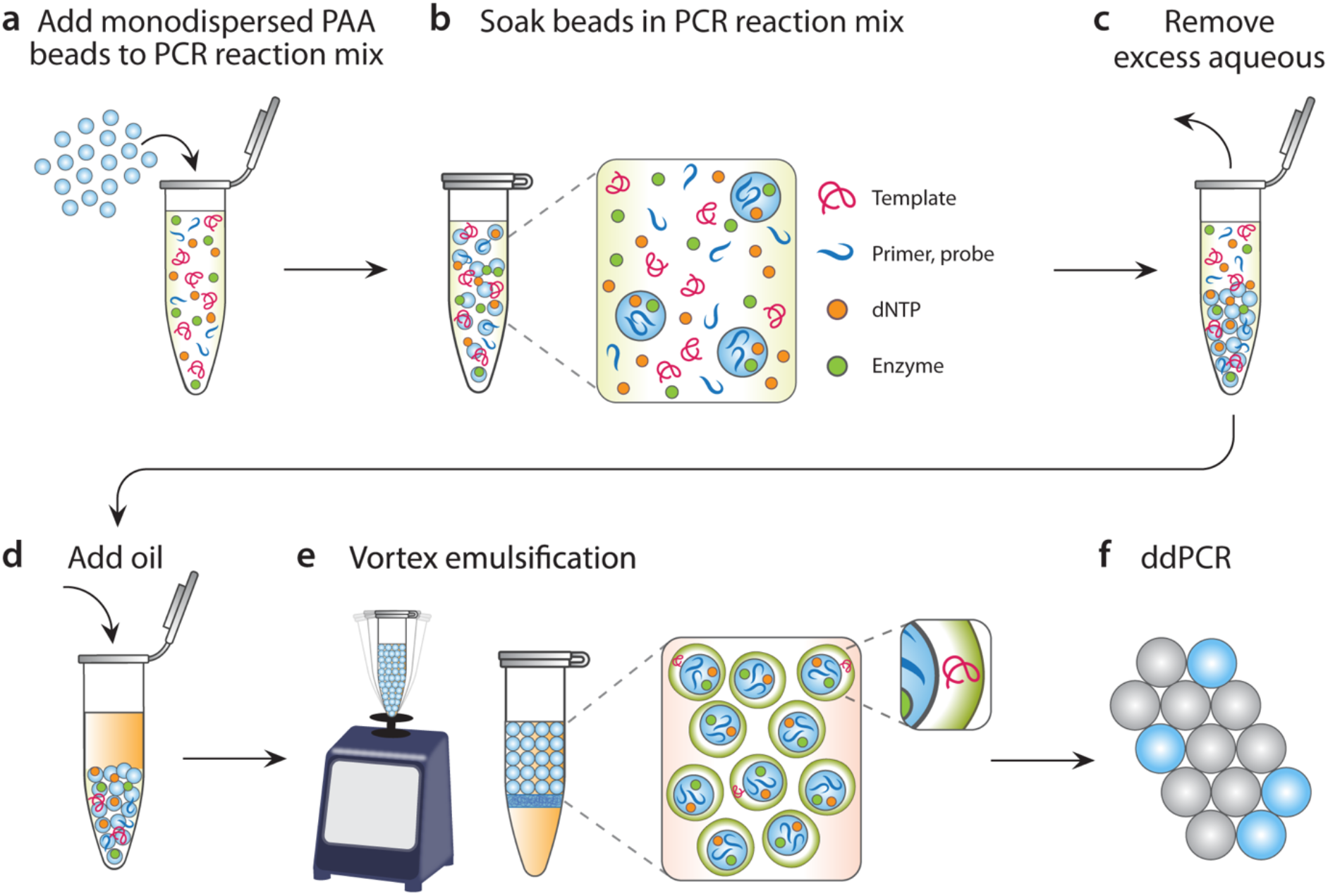
Schematic of PTE workflow. Monodispersed polyacrylamide (PAA) beads are added to the PCR reaction mix (**a**) and soaked with enzymes and molecules (**b**). After removing excess aqueous phase (**c**), oil with stabilizing surfactant is introduced (**d**) and the mixture is uniformly emulsified by vortexing (**e**). For digital droplet PCR, target sequences are encapsulated and amplified in the droplets, generating a detectable fluorescence signal (**f**).

PTE encapsulates reagents in the initial sample into droplets, allowing compartmentalized reactions like what is normally achieved with microfluidics. Compounds smaller than the hydrogel pore size are absorbed before emulsification and are thus present in the final droplet containing the hydrogel. Larger entities end up in the thin layer of aqueous solution surrounding the hydrogel, as illustrated in Fig. 1e. We use vortexing for PTE due to its reproducibility, although other agitation techniques are compatible, like pipetting and tube flicking.

### Optimizing PTE

Vortexed emulsions are simple to make but polydispersed and of limited value for precision biology (Fig. 2a), while microfluidic emulsions require specialized devices and skill, but exhibit superior monodispersity and are of utmost value (Fig. 2a). An optimal method for sample encapsulation would, thus, combine the simplicity of vortexing with the quality of microfluidics. PTE accomplishes this by exploiting the rigidity of particles to resist droplet breakup below the particle size, even with vortexing. Nevertheless, the time to reach the final droplet size and the monodispersity of the resultant emulsion still depend on fluid and particle properties. For example, particle properties like size and self-affinity, and solution properties like viscosity and interfacial tension, affect the droplet templating process. To characterize the impact of these parameters on emulsion monodispersity, we perform PTE with different hydrogel materials and solution interfacial tensions (Fig. 2b). We fix particle size, carrier oil, and oil-soluble surfactant, since they are usually dictated by the needs of the biological reactions, and inflexible.

To modulate interfacial tension, we add aqueous-soluble surfactants to the droplet phase that are compatible with most biochemical reactions. When the aqueous-phase surfactant is omitted, the vortexed droplets are polydispersed for all hydrogel types (Fig. 2b); this may be due to inter-particle affinity and high water-oil interfacial tension, preventing breakup of large, non-uniform, multi-core droplets. Increasing vortexing up to 20 min does not appreciably change emulsion monodispersity (Supplementary Fig.S1a). By contrast, when we include surfactants in the aqueous phase, particle affinity and interfacial tension are reduced; monodispersed single-core droplets are generated in 30 sec of vortexing. Droplets formed from polyacrylamide (PAA) and polyethylene glycol methacrylate (PEG) particles yield the best emulsions (Fig. 2b). Agarose-particle emulsions are less uniform, possibly due to their self-affinity. Other aqueous-phase surfactants may also be used, although the requisite vortexing parameters and emulsion monodispersity may differ (Supplementary Fig. S1b, c). We have focused on ~50 *μ*m hydrogels, but other particle types are compatible with the method, including different sizes, hydrogel chemistries, and porosities. While we focus on fluorinated oil and surfactant for our carrier phase, because they are predominant in the field due to their value for cell and molecular biology, other formulations are compatible, including silicon and hydrocarbon oils and surfactants. Other polar phases can also be used, provided they form stable emulsions, which should be valuable for generating core-shell structures. These properties should provide much needed flexibility for extending PTE to other areas, such as cost-effective and scalable encapsulation of compounds in double emulsions.

**Figure 2.**
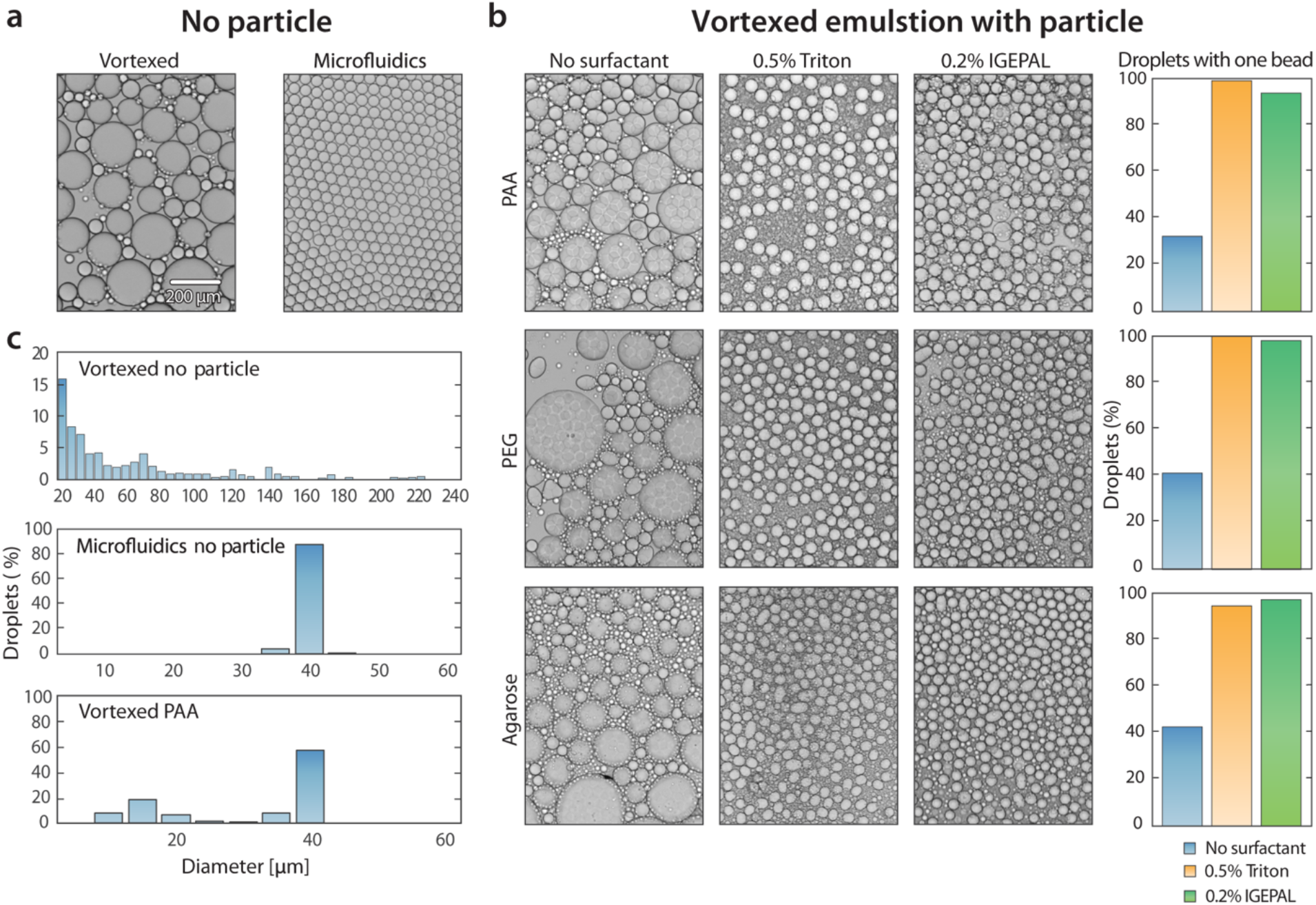
Size uniformity of droplets generated by PTE. (**a**) While microfluidically-prepared emulsions are monodispersed, untemplated vortexed emulsions are not. (**b**) For certain conditions, vortexing with particle templates allows generation of monodispersed emulsions. To identify the best conditions, we test different particle compositions and aqueous-soluble surfactants. The fraction of droplets containing a single hydrogel particle is determined by image analysis (bar graphs on right). PAA and PEG-templated emulsions with Triton yield 98.8% (n= 433) and 99.1% (n= 683) singleparticle droplets, respectively. (**c**) We quantify the monodispersity of the resultant emulsions by image analysis, plotting as histograms. Untemplated vortexing yields polydispersed emulsions, with 4.3% of diameters between 35–40 μm (n= 561), while microfluidic generation yields monodispersed emulsions, with 95.7% of droplet diameters between 35-38 μm (n= 816). PTE droplets contain either a single hydrogel core (58%, 35-40 μm) or no hydrogel particle (42%, < 35 μm). The no-particle droplets, however, due to their small sizes, contribute to < 5% of the total sample volume (n= 1421).

In addition to the desired monodispersed droplets with size similar to the templating particles, PTE generates tiny “satellite droplets” containing no particles. The number of satellites depends on the amount of excess aqueous solution surrounding the particles, emulsion interfacial tension, and vortexing time and power. To reduce their number, excess aqueous solution should be removed from the particle-sample mixture prior to vortexing. Nevertheless, while aesthetically unpleasing, satellites contribute negligibly to biological reactions performed in the emulsions, because they usually comprise a relatively small fraction of the total sample volume (<3%).

Engulfed volume can be predictable by considering vortex power, surface tension and particle size and so on. Vortexing, however, generate a distribution of velocities in the sample, and each droplet experiences a random sample of these velocities during emulsification and so this is only a rough estimation. Nevertheless, vortexing is a reasonably controlled method for agitating fluids and, thus, it is possible to identify a power that yields mostly single core droplets with uniform engulfment volumes, as we have shown.

A common challenge in droplet microfluidics is the efficient encapsulation of discrete entities, like beads and cells. Microfluidic techniques normally encapsulate discrete entities randomly, resulting in inefficient Poisson loading in which only a small fraction of the droplets are properly loaded. A unique and valuable property of PTE is that every droplet of the appropriate size contains one templating particle (Fig. 2b). If these particles are an essential component of the reaction, most droplets will contain one. Other components, however, such as cells, beads, and DNA molecules, are loaded randomly. Indeed, efficient hydrogel encapsulation is a key step in recently reported single cell sequencing technologies and exploited in commercial instruments^24–26^.

### PTE allows accurate DNA quantitation with digital droplet PCR

A notable commercial success of droplet microfluidics is ddPCR, an alternative to quantitative PCR for nucleic acid analysis that is more sensitive, accurate, and provides absolute concentration measurements ^18,27,28^. In the approach, the nucleic acids to be quantitated are microfluidically encapsulated into millions of aqueous droplets at limiting dilution, and subjected to PCR. During amplification, droplets containing targets become fluorescent while those devoid remain dim, enabling the absolute quantification of nucleic acid concentration by counting fluorescent droplets. While valuable, all described ddPCR approaches require single-use microfluidic chips, costly control hardware, and custom reagents. A method for performing ddPCR without microfluidics and using commonly available reagents would, thus, be beneficial to many labs.

PTE allows facile, microfluidic-free ddPCR.To illustrate this, we encapsulate several DNA samples at different concentrations of a target molecule (Fig. 3a). Just as in microfluidic ddPCR, increasing target concentration increases the number of fluorescent droplets. To determine whether this allows concentration estimation, we follow conventional ddPCR analysis and quantify droplet fluorescence, plotting the results as fluorescence versus diameter (Fig. 3b). Three droplet populations are visible, at low fluorescence and small diameter (satellites), at the expected 30-40 *μ*m diameter and low fluorescence (PCR-negative), and similar size range but high fluorescence (PCR-positive). We ignore the satellites and model target concentration for the correctly-sized droplets via Poisson statistics,

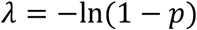

where *λ* is the template copy number per droplet and *p* is the positive fraction. The measured concentration follows the expected scaling over the three-decade tested range (Fig. 3c). This shows that PTE ddPCR performs like microfluidic ddPCR.

**Figure 3.**
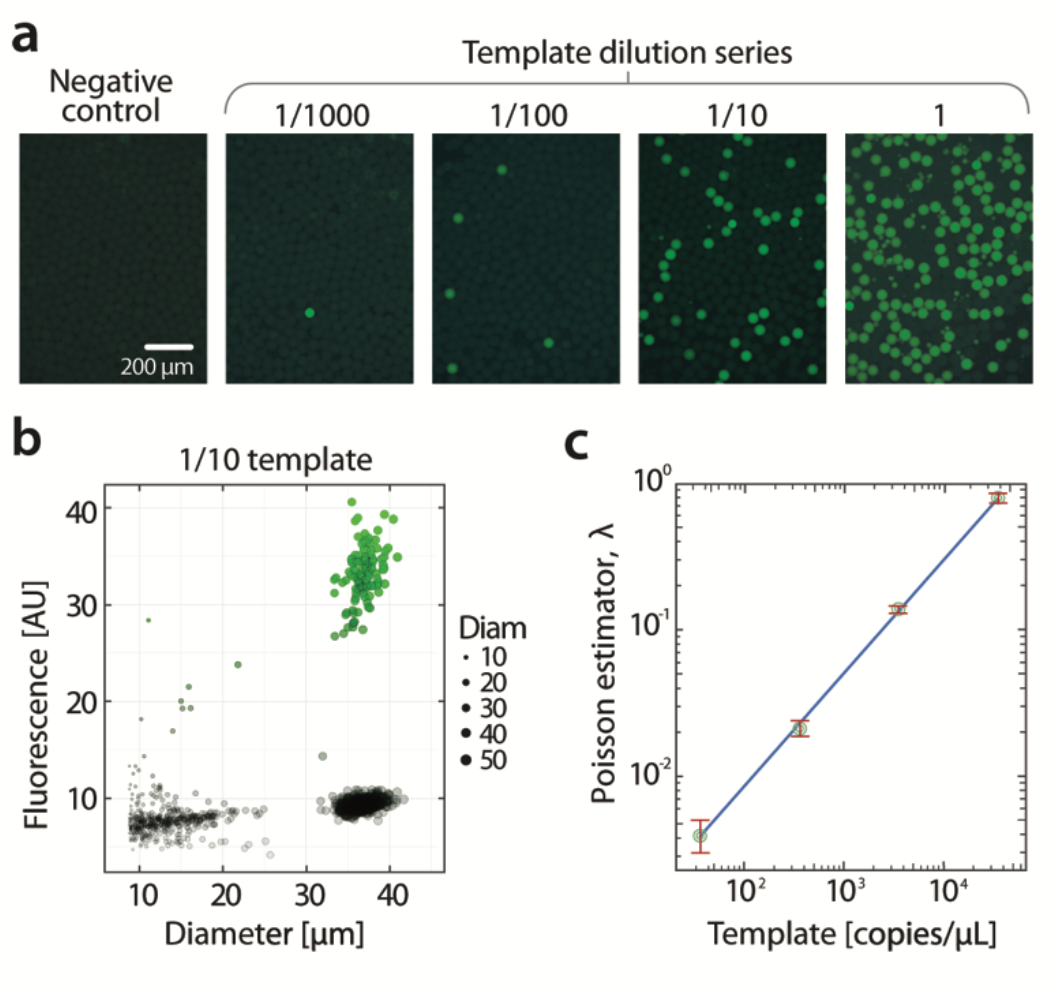
ddPCR assay with droplets prepared by PTE. (**a**) Fluorescence images of droplets after PCR amplification with TaqMan probes and primers for yeast genomic DNA templates at varying dilution factors. The fractions of observed fluorescence-positive droplets correspond with the template concentrations. (**b**) Scatter plot showing the size and fluorescence distribution from a sample in the dilution series. The population with low fluorescence (< 20 AU) and small diameter (< 30 μm) is composed of droplets containing no hydrogel particles (bottom-left). The population with expected diameter (30-40 μm) consists of single-hydrogel-core droplets. They form two tight clusters: high fluorescence (PCR-positive) and low fluorescence (PCR-negative, no-template droplets). (**c**) The average template copy number per droplet estimated by assuming a Poisson distribution scales with the controlled template concentrations over the tested three-decade range (R^2^= 0.9994 and error bar depicts standard error).

Essential to the PTE method are the hydrogel particles that template the droplets, which we have used microfluidics to make. Even with microfluidically-made particles, PTE represents a substantial simplification for ddPCR, since one large batch of synthesized particles can be used for many analyses. Nevertheless, the use of microfluidics undercuts the main advantage of PTE for labs entirely lacking this expertise. An optimal implementation of PTE would obviate all microfluidics and, ideally, use only commercially available components.

Indeed, hydrogel microspheres with a variety of compositions, sizes, and uniformity can be purchased from commercial vendors. These spheres are usually sold as components for purification columns and, thus, quality-controlled and free of contaminants that could interfere with reactions. To show that PTE can be performed with commercial hydrogels, we purchase quasi-monodispersed PAA spheres, ranging from 45-90 *μ*m diameter; this size distribution is larger than for particles made with microfluidics, which are typically below 5%, but is acceptable for most applications, including ddPCR. To demonstrate this, we use the particles to perform ddPCR with PTE, and observe similar droplet fluorescence properties (Fig. 4a). The larger diameter distribution results in broader scatter in the plot, both in size and fluorescence, but the PCR positive and negative populations are nevertheless clearly discernable (Fig. 4b). We thus vary target concentration and perform standard ddPCR analysis, achieving accurate measurements over the same range (Fig. 4c). When the variability in droplet size is included using multiple Poisson distributions weighted by droplet volume, the correction factors for estimated copy numbers are small, ranging from 0.1% (for the lowest concentration) to 4.5% (for the highest concentration).

**Figure 4.**
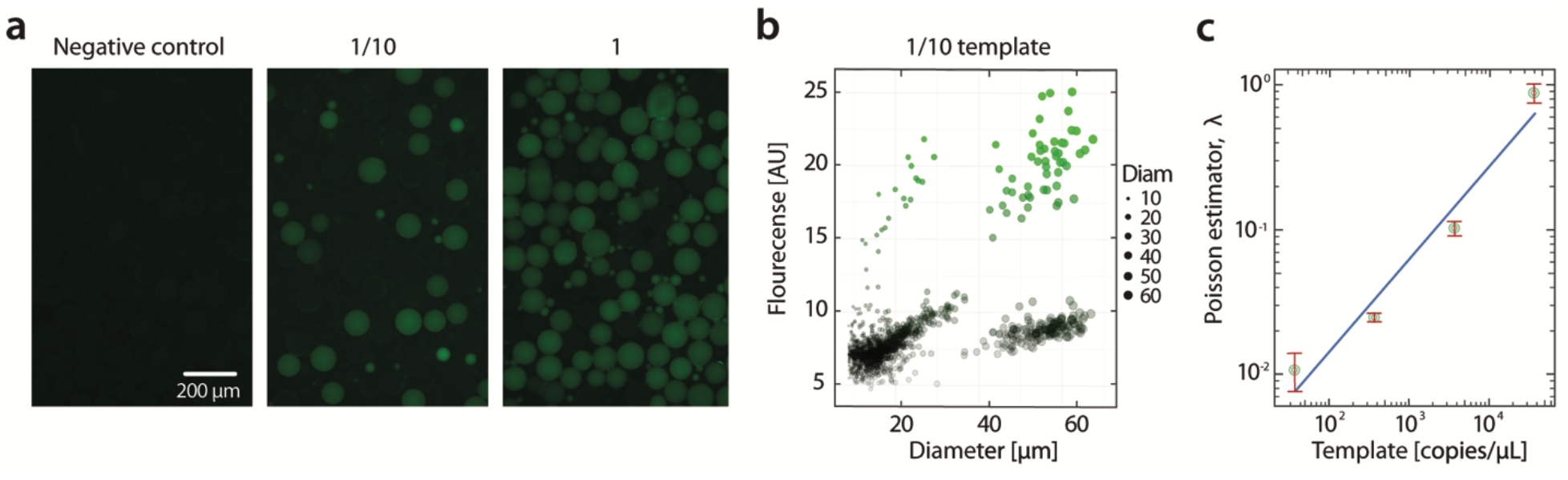
Demonstration of ddPCR quantitation with commercially-available hydrogel particles. (**a**) Fluorescence images of droplets after PCR amplification of yeast genomic DNA at different concentrations. (**b**) Scatter plots show that the hydrogel-templated droplets are less uniform in size. But fluorescence-positives and negatives are clearly distinguishable from each other, enabling quantitation by image analysis. (**c**) The Poisson estimator values obtained by using multiple Poisson distributions weighted by droplet volumes show a linear correlation with the template concentration (R^2^= 0.9409 and error bar depicts standard error).

Access to monodispersed particles is critical for PTE, which will in most cases be the principal barrier to its implementation. We use microfluidically-made particles to characterize PTE, since they are monodispersed and thus allow accurate measurement of droplet volume variation. However, as we have shown, commercially available beads that are relatively uniform will suffice for many applications.

### Multiplexed PTE-ddPCR

Like other PCR analysis methods, ddPCR can be multiplexed using probes labeled with different fluorescent dyes. Since ddPCR acts on molecules in droplets, this provides unique measurement opportunities not possible with common “bulk” methods, like the physical association of distinct sequences. This is valuable for a variety of important applications in genomic biology, including characterizing virus diversity, phasing microbial genomes, haplotyping cancer genomes, measuring mRNA splice forms, and characterizing length distributions of target molecules in solution ^29–31^. To demonstrate that PTE-ddPCR can be multiplexed, we use it to analyze a mixture of lambda virus and yeast (*S. cerevisiae*) genomic DNA. We use TaqMan probes targeting either the lambda virus (red) or yeast (green) genomes. The DNA of both organisms is mixed together, and the sample emulsified with PTE. We find that, as expected, many droplets are pure red or green, indicating that they contain either lambda or yeast genomic DNA, respectively (Fig. 5a). However, in rare instances, a droplet contains one of each target and, thus, is double-positive, appearing yellow (Fig. 5a, merged). Since these nucleic acids do not physically associate, the likelihood of a double positive is described by Poisson double-encapsulation ^32^. Deviations from Poisson statistics thus represent associations of sequences ^31^.

**Figure 5.**
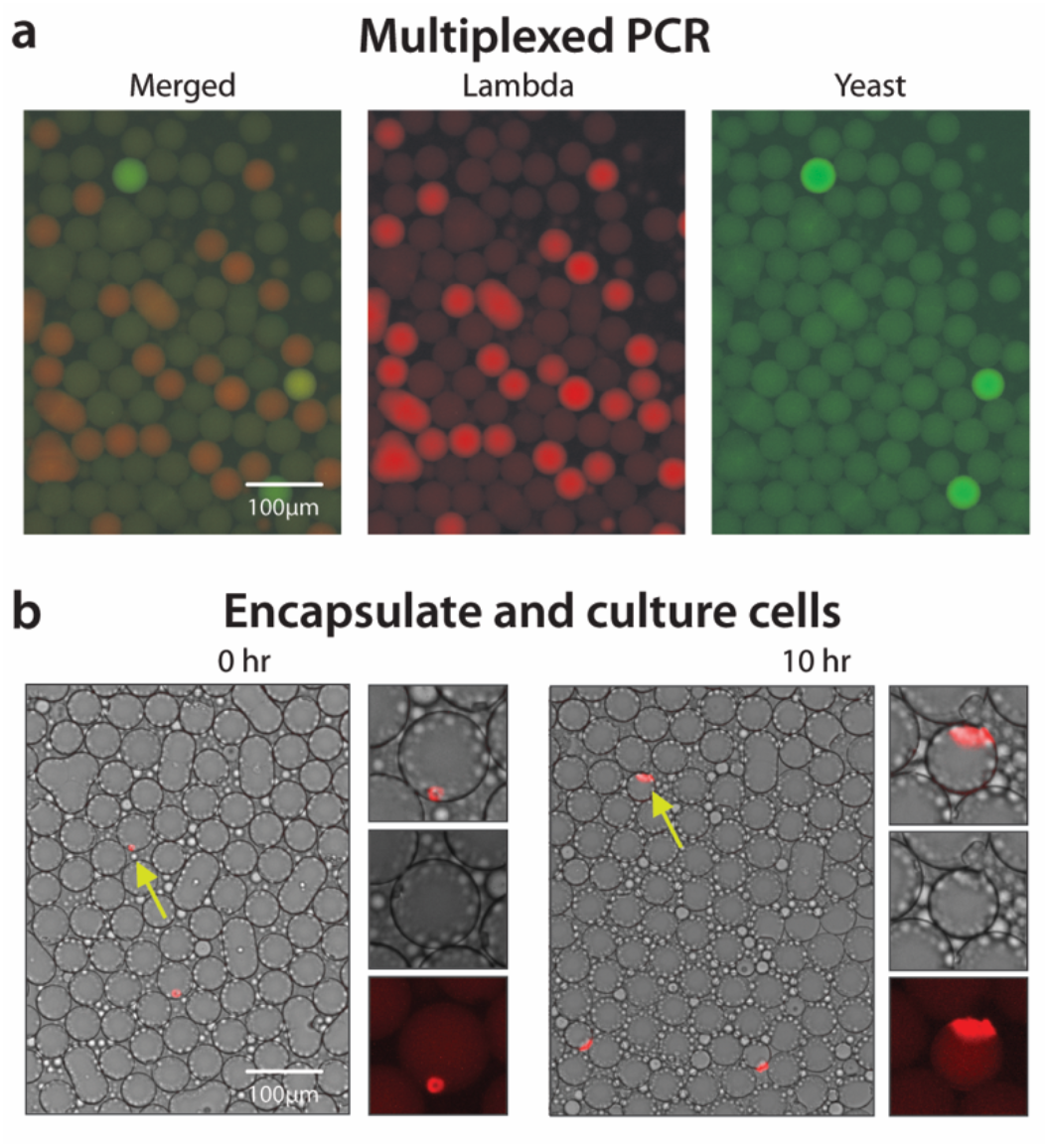
PTE for multiplexed ddPCR and cell culture. (**a**) Probes targeting lambda virus or yeast are fluorescently labeled with Cy5 (red) and FAM (green), respectively. (**b**) Yeast cells grow in droplets prepared by PTE. After 10 hours of incubation, colonies grown from single encapsulated cells can be detected by endogenous YFP fluorescence.

### PTE for single cell biology

Single cell biology is growing in importance, fueled by microfluidic tools for encapsulating, culturing, and analyzing huge numbers of single cells ^9,33^. New methods for ultrahigh-throughput single cell sequencing are valuable for studying the importance of heterogeneity to phenotypes and disease. These methods, however, require co-encapsulation of a cell and barcoded bead in every droplet, an inefficient process that requires specialized microfluidic chips and control systems. Similar encapsulations can be efficiently achieved with PTE much more simply, since it already uses hydrogel spheres to template droplet generation. To demonstrate this, we use PTE to encapsulate single yeast cells. The PAA hydrogels are added to a suspension of yeast cells and the mixture emulsified by vortexing. The cells are suspended at low concentration so that most droplets are empty, but a small fraction contain single cells, just as in microfluidic methods. Because the micron-scale yeast cannot diffuse into the nanometer hydrogel pores, they end up in the aqueous shell near the periphery of the droplets (Fig. 5b). The number of yeast encapsulated per droplet can be controlled with cell concentration in the original sample.

The droplet environments are compatible with yeast growth, because PAA is a biologically inert hydrogel that comprises >95% aqueous solution. Consequently, when we incubate the emulsions, the yeast cells grow into clonal microcolonies (Fig. 5b). Such microcolonies are useful for microbe cultivation, sequencing, and metabolite screening ^9,34,35^.

We describe particle-templated emulsification (PTE), an approach for generating compartmentalized reactions in monodispersed droplets with vortexing. The particles afford an independent and avenue for introducing compounds into the droplets, for example, by functionalizing with reactive groups, oligos, and proteins ^36,37^. Reversible cross linkers can release functional groups or melt gels after encapsulation, providing a fully-liquid environment that might have advantageous chemical kinetics. For example, recently described single cell RNA and DNA sequencing require the encapsulation of unique “barcode” sequences in droplets, which can be achieved by functionalizing each bead with a unique sequence using split-pool techniques ^20^. Pre-functionalized beads can be used in PTE to generate and encapsulate cells in the emulsion, and introduce the barcode sequence. This yields an emulsion equivalent to what is currently generated with microfluidics at a fraction of the cost and microfluidics-free, making adoption of these methods simpler and more affordable.

PTE is gentle on biological samples and uses biocompatible chemicals and particles, making it applicable to many reactions. For example, recently described digital droplet multiple displacement amplification techniques have demonstrated compartmentalized amplification for quantitative sequencing of single cell and low-input samples; however, all require microfluidics or custom hardware. PTE can perform this and many other compartmentalized reactions. Moreover, PTE is more scalable than conventional microfluidic emulsification because the time to emulsify a sample does not increase appreciably with sample volume. This should make it valuable for implementing droplet encapsulation into high throughput workflows performed in well plates, which can contain hundreds of separate samples that each need to be emulsified. With PTE, particles, oil, and surfactant can be added to all wells, and the plate agitated using plate-vortexers or pipetting to generate the emulsions. Combined, these properties should make PTE especially valuable for high throughput sequencing workflows involving robotic preparation of samples that are otherwise difficult to use with microfluidics.

While PTE prepares huge numbers of droplet assays without special equipment, many applications require analyzing or sorting droplets, like the ddPCR method we have demonstrated here ^27^, or enzyme evolution methods described previously ^38,39^. Microscopy, which is widely available in the biological laboratory can be used for simple droplet analysis, as we have shown. Specific droplets can even be recovered manually with a pipette. In some cases, however, microscopy and manual picking may not suffice, and higher throughput analysis will be needed. In these cases, the best option may be flow cytometry, in which the PTE emulsion can be re-encapsulated in an aqueous carrier using *in vitro* compartmentalization techniques ^40–42^. The resultant water-suspended droplets can be directly analyzed and sorted with flow cytometry ^43^. Alternatively, chemical techniques can exploit the hydrogel scaffold, attaching, for example, fluorescent reaction products to the gel backbone. The solid particles can be re-dispersed in aqueous solution and subjected to flow cytometry ^22^.

## Conclusions

Droplet microfluidics has enabled ultrahigh-throughput digital biology, to precision-analyze large numbers of individual molecules and cells. Uniformity of the droplet compartments is crucial, but necessitates microfluidic devices and expertise, limiting adoption by the community at large. Here, we report a novel method to prepare monodispersed emulsions by vortexing that obviates microfluidics. PTE generates droplets with uniformity like microfluidics and is useful for similar applications. In addition, it can encapsulate reagents in the droplets, including beads, cells, and molecules, and is compatible with biological operations like cell culture and ddPCR. This simple but fundamental method should facilitate adoption of droplet compartmentalization for biological applications and advance the field of digital biology.

## Acknowledgments

We thank Leqian Liu for helpful discussions and David Sukovich for primer and probe sets. This work was supported by the National Science Foundation through a CAREER Award (grant number DBI-1253293); the National Institutes of Health (NIH) (grant numbers R01-EB019453-01, DP2-AR068129-01, R01-HG008978).

